# Efficiency, accuracy and robustness of probability generating function based parameter inference method for stochastic biochemical reactions

**DOI:** 10.64898/2026.01.21.700833

**Authors:** Shiyue Li, Yiling Wang, Zhanpeng Shu, Ramon Grima, Qingchao Jiang, Zhixing Cao

**Affiliations:** State Laboratory of Bioreactor Engineering, East China University of Science and Technology, Shanghai 200237, China; Department of Chemical Engineering, Queen’s University, ON K7L 3N6, Canada; College of Electrical Engineering, Shanghai Dianji University, Shanghai 201306, China; School of Biological Sciences, University of Edinburgh, Edinburgh, EH9 3BF, United Kingdom

**Keywords:** Probability generating function, parameter inference, stochastic modeling, model selection

## Abstract

Biochemical reactions are inherently stochastic, with their kinetics commonly described by chemical master equations (CMEs). However, the discrete nature of molecular states renders likelihood-based parameter inference from CMEs computationally intensive. Here, we introduce an inference method that leverages analytical solutions in the probability generating function (PGF) space and systematically evaluate its efficiency, accuracy, and robustness. Across both steady-state and time-resolved count data, our numerical experiments demonstrate that the PGF-based method consistently outperforms existing approaches in terms of both computational efficiency and inference accuracy, even under data contamination. These favorable properties further enable the extension of the PGF-based framework to model selection—a task typically considered computationally prohibitive. Using time-resolved data, we show that the method can correctly identify complex gene expression models with more than three gene states, a task that cannot be reliably achieved using steady-state data alone.

## I. Introduction

Biochemical reactions are inherently stochastic, arising from the random collisions of biomolecules, whose movements are naturally unpredictable. Gene expression is a quintessential example of this phenomenon, with extensive experimental evidence confirming its stochasticity [1], [2], [3], [4], [5]. For clarity, we will primarily use gene expression to illustrate our proposed method, though the approach is generalizable. The stochastic nature of these reactions necessitates a probabilistic framework for quantitative kinetic analysis, enabling a more precise understanding of molecular-level processes [6], [7].

A biochemical reaction system can be generally represented by a set of reaction equations [8], [9]:

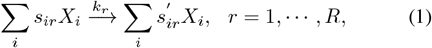

where *s*_*ir*_ and 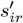 are the stoichiometric coefficients of species *X*_*i*_ in reaction *r*. Assuming the law of mass action, the rate of reaction *r* is given by

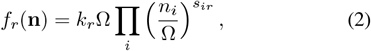

where *k*_*r*_ is the rate constant, **n** = [*n*_1_,, *n*_*N*_]^⊤^, *n*_*i*_ is the molecule count of species *X*_*i*_, and Ω is the reaction volume. A fundamental task in analyzing the kinetics of the reaction system in Eq. (1) is inferring the kinetic parameters *k*_*r*_ from observed molecule counts (*n*_*i*_) ∈ 𝕀 of certain species—a process known as parameter inference or estimation in systems biology [10], [11], [12], or system identification in control theory [13], [14]. 𝕀 is the set of natural numbers.

Parameter inference is fundamentally an inverse problem that necessitates repeated forward computations of the kinetic model. Given the various approaches available for kinetic model computation, the inference methods in the literature can be broadly classified into four groups. **The first group** employs maximum likelihood estimation (MLE) combined with finite state projection (FSP) [11], [15], [16], [17]. FSP solves a set of chemical master equations (CMEs) [9], [18], which are difference-differential equations commonly used to describe stochastic reaction kinetics. This approach assumes that the probability of molecule counts exceeding a certain threshold (truncated size) is zero [19], [20]. However, the computational efficiency of these methods declines rapidly as the number of species, and consequently the number of equations, increases exponentially. Moreover, the selection of the truncated size requires careful consideration to achieve an intricate balance between computation load and precision. **The second group** employs the method of moments (MOM), where a few low-order moments are calculated both from the molecule count data and the kinetic models, and then used to generate a Gaussian-like synthetic likelihood for inference [12], [21], [22], [23], [24]. These methods are computationally efficient, requiring the solution of only a few differential equations. However, their accuracy can be unsatisfactory, especially when higher-order moments are needed to derive a sufficient number of moment equations for inference. In such cases, the accuracy of moments computed from small sample sizes can be compromised [10]. Additionally, if more than one species participates in a reaction, i.e., ∑_*i*_ *s*_*ir*_ *>* 1, the moment equations derived from the corresponding CMEs are not closed, necessitating the use of various moment closure methods [18], [25], [26]. Moment closure is inherently an approximation, potentially introducing another layer of inaccuracy. **The third group** employs an Approximate Bayesian Computation (ABC) scheme combined with the Stochastic Simulation Algorithm (SSA) for parameter inference [27], [28], [29]. ABC approximates the posterior distribution by simulating data under various parameter values and comparing it to observed data. Parameters yielding simulations that closely match the observed data are accepted as approximations of the true posterior. This approach is advantageous as it bypasses explicit likelihood calculations, with SSA providing an exact method for generating simulation data. However, this framework has drawbacks, including the need for large simulation samples to accurately approximate the posterior, which can be computationally expensive, and sensitivity to tuning parameters such as the tolerance level and distance metric.

The final group is the PGF-based inference method [30], [31], [32], which we systematically investigate in this work. This method computes the empirical PGF directly from count data and compares it with the analytical PGF solution derived from the model, using either the density power divergence [30], [31] or the mean squared error [32] as the objective function. Minimizing this discrepancy yields the inferred kinetic parameters. Ref. [32] has demonstrated several advantages of the PGF-based inference method: (i) Analytical PGF solutions are available for a broad class of gene expression models. Traditionally, these solutions have been used by performing Taylor expansions to recover probability mass functions, followed by maximum likelihood estimation (MLE) for parameter inference. However, this approach is numerically demanding—particularly because PGF solutions often involve hypergeometric functions that require high-order derivatives, which are computationally unstable and require high numerical precision. As a result, such methods are not widely adopted [33], [34]. In contrast, the PGF-based method circumvents the need for differentiation by directly evaluating the PGF over a range of variable values, thereby improving both stability and computational efficiency. This approach enables full utilization of existing PGF solutions. (ii) The PGF-based method achieves computational efficiency comparable to MOM, while maintaining inference accuracy on par with MLE. Building on these advantages, we systematically evaluate the accuracy, efficiency, and robustness of the PGF-based method under two types of data contamination: binomial downsampling and outliers. Furthermore, we extend the PGF-based framework in Ref. [32] from steady-state to time-resolved count data. Within this extended setting, we develop a model selection strategy based on cross-validation. Using this approach, we demonstrate that time-resolved data enables reliable identification of complex gene expression models with more than three gene states—a task that cannot be accomplished using steady-state data alone.

The paper is organized as follows. Section II introduces the preliminaries of PGF theory and reviews analytical solutions for representative gene expression models. Section III presents the PGF-based inference method for steady-state count data. Section IV evaluates its computational efficiency, accuracy, and robustness under various conditions. Section V extends the method to time-resolved count data and develops a model selection framework based on PGF inference. Section VI concludes the paper and discusses future research directions.

## II. Preliminary

Consider a reaction system consisting of *N* species (*X*_*i*_ for *i* = 1, *· · ·, N*) and *R* reactions as defined by Eq. (1) with reaction rates given by Eq. (2). The kinetics of this system can be effectively described using the probabilistic framework of CMEs

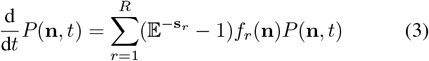

Where *P* (**n**, *t*) represents the probability of observing *n*_*i*_ copies of molecule *X*_*i*_ for *i* = 1, *· · ·, N* in the system at time *t*. The vector **s**_*r*_ is defined as

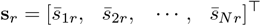

with 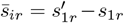. The step operator 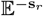 acts on a general function *f* (*n*_1_, · · ·, *n*_*N*_) as follows

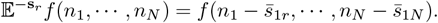

This indicates that applying the operator shifts the arguments of the function *f* by subtracting the corresponding components of the vector **s**_*r*_. Solving Eq. (3) is challenging due to the presence of both discrete variables (*n*_*i*_, which are integers) and continuous variables (*t*). The PGF method offers a way to circumvent this challenge. The PGF is defined as

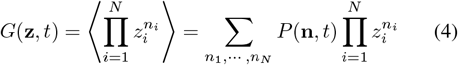

in which **z** = [*z*_1_,, *z*_*N*_]^⊤^ and ⟨·⟩ is the expectation operator. Essentially, PGF is a multi-dimensional *z*-transform of the function *P* (**n**, *t*).

By applying Eq. (4), Eq. (3) can be conveniently transformed into a set of partial differential equations (PDEs). These resulting PDEs can then be tackled using various standard methods for solving PDEs. This approach, known as the PGF method, has been effectively employed to solve a wide range of kinetic models, as summarized in Table I. In the following sections, we will introduce some properties of the PGF, which allow the construction of the PGF for more complex systems by using the solutions in Table I as foundational building blocks.

**TABLE I.**
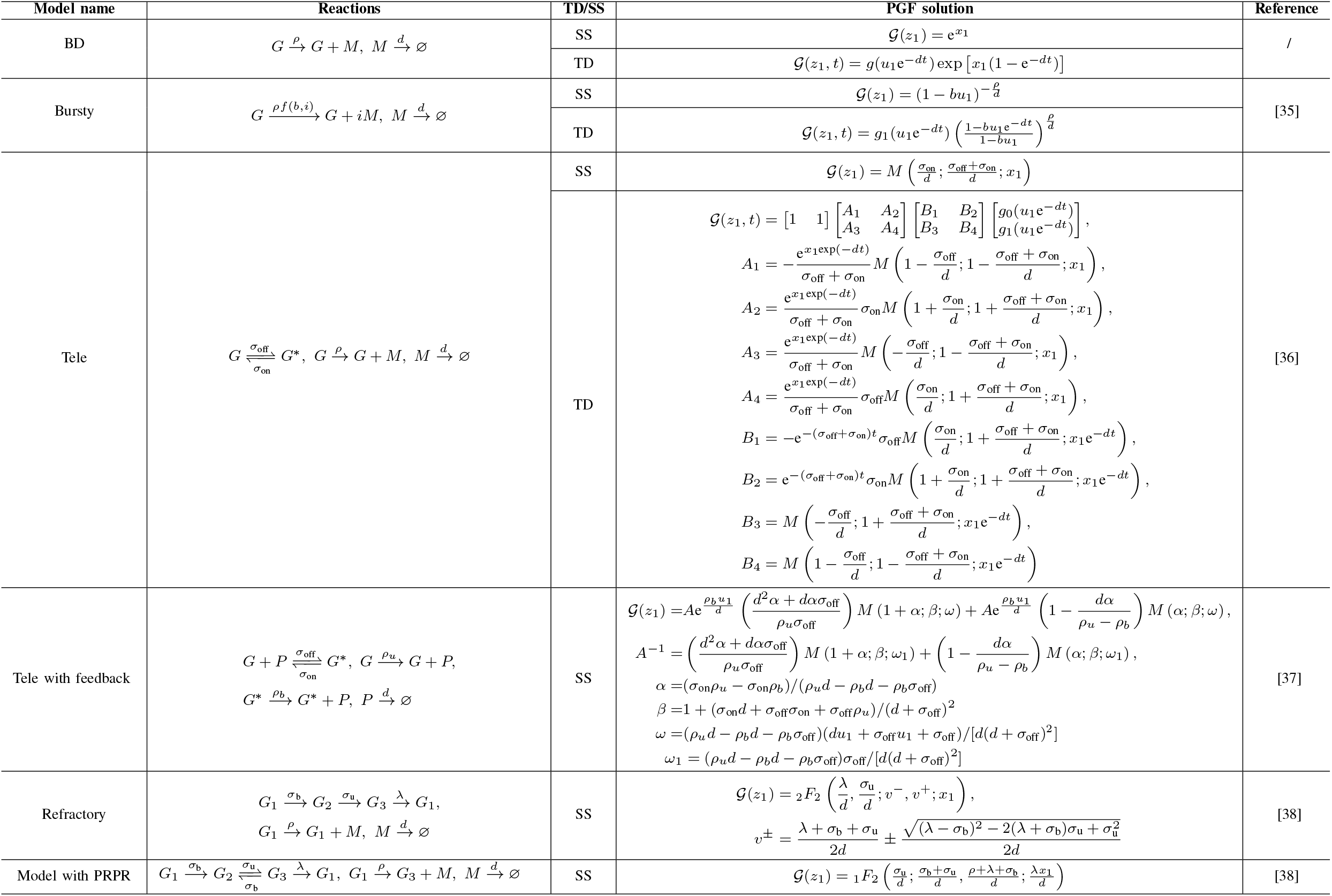
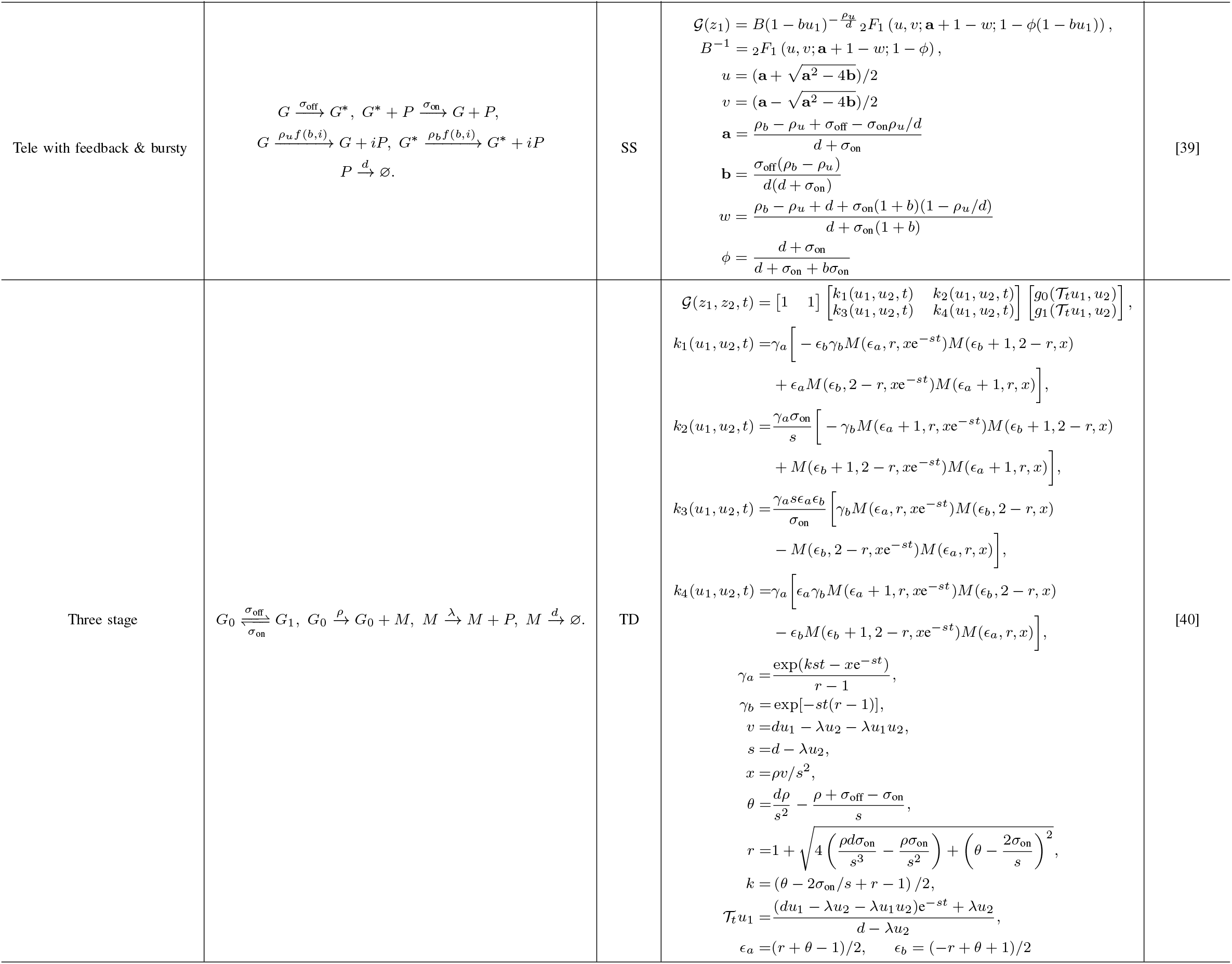
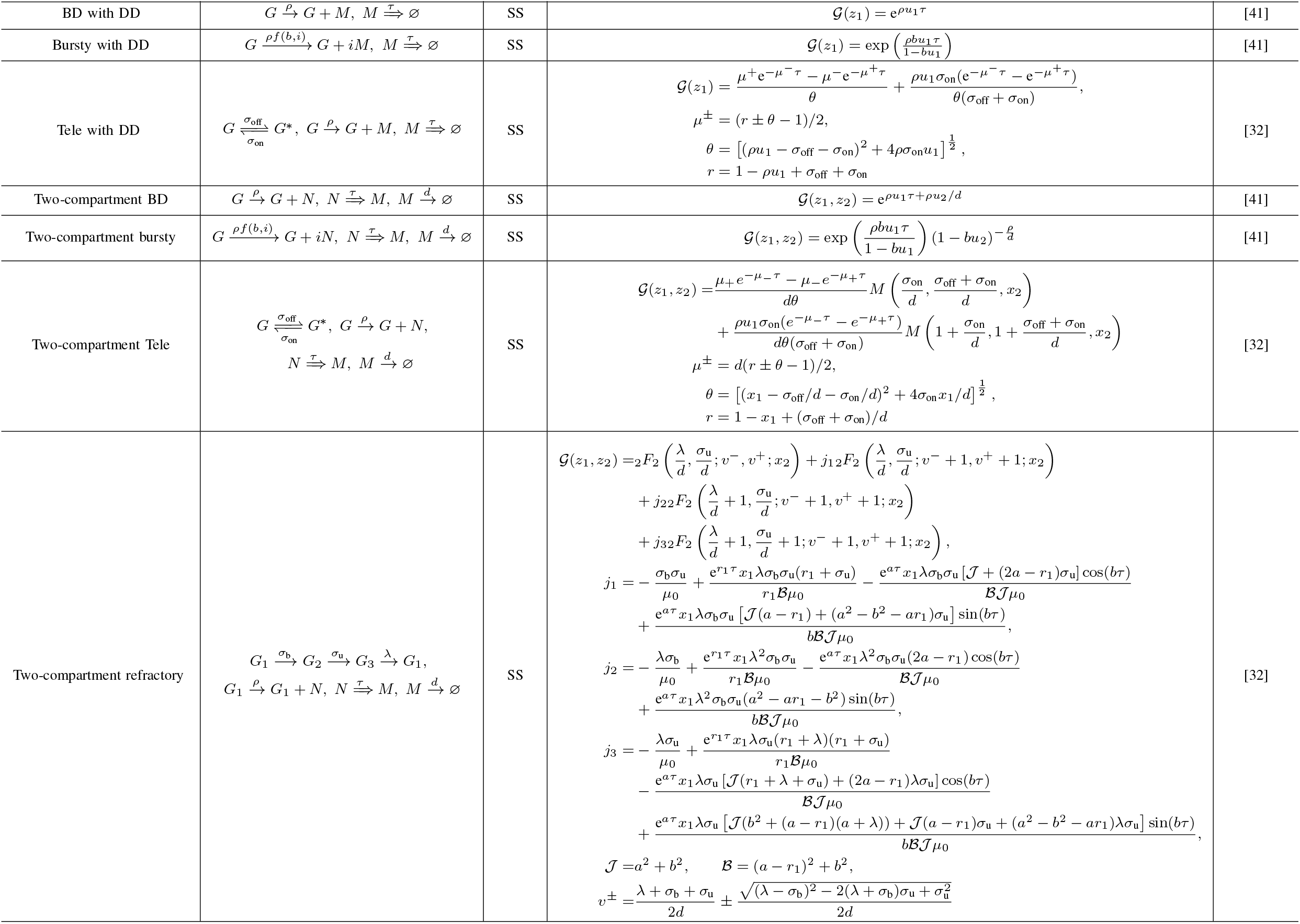
Zoo of PGF solutions: TD stands for time-dependent, SS stands for steady-state, BD stands for birth-death, Tele stands for telegraph, PRPR stands for polymerase recruitment and pause release, and DD stands for delayed degradation. *f* (*b, i*) = *b*^*i*^*/*(1 + *b*)^*i*+1^, *u*_1,2_ = *z*_1,2_ −1, *x*_1,2_ = *ρu*_1,2_*/d, M* (.) stands for kummer confluent hypergeometric function

### (P1) Binomial partitioning

Given that each random variable *m*_*i*_ in the vector **m** is independently drawn from the corresponding random variable *n*_*i*_ in the vector **n** with a binomial distribution, the probability mass function for *m*_*i*_ given *n*_*i*_ is:

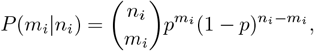

where *p* is the success probability in the binomial distribution. The PGF 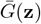 for the random vector **m** can be derived using the relationship between the PGFs of **n** and **m**. The PGF of **m** is given by:

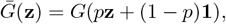

where **1** = [1, *· · ·*, 1]^⊤^. This transformation reflects the fact that for a binomial distribution, the generating function of *m*_*i*_ given *n*_*i*_ is 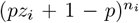. Applying this transformation to each component of **z** in the original PGF yields the PGF for the vector **m**. See the supplementary information of Ref. [42] for a detailed proof. This process closely resembles the partitioning of molecules between daughter cells during cell division, where each molecule in the mother cell has a certain probability of being allocated to one daughter cell.

### (P2) Marginalization

The PGF marginalized with respect to the random variable *n*_*i*_ is obtained by setting *z*_*i*_ = 1 in the PGF. Thus, the marginalized PGF is expressed as 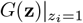. This operation effectively removes the dependence on the random variable *n*_*i*_, resulting in the PGF for the distribution of the remaining variables.

### (P3) Summation of all variables

Given that *g*(*z*) is the PGF for the random variable 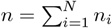, the PGF *g*(*z*) is given by

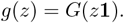

This expression indicates that the PGF of the sum of the random variables *n*_*i*_ is obtained by evaluating the original PGF *G*(**z**) at the point where each component *z*_*i*_ of the vector **z** is replaced by the same value *z*. This can be seen from

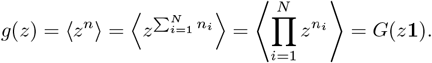

### (P4) Independence

Given two random vectors **n, m ∈** 𝕀^*N*^, with their respective PGFs *G*_1_(**z**_1_) and *G*_2_(**z**_2_), if each random variable *n*_*i*_ in **n** is independent of *m*_*j*_ in **m** for any *i* and *j*, the joint PGF of **n** and **m** is given by

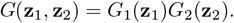

This means that the joint PGF of two independent random vectors is simply the product of their individual PGFs. Consider a new random vector **q** = **n**+**m**. Its corresponding PGF, 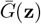, can be directly obtained by applying properties P1 and P4, resulting in the following expression:

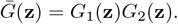

### (P5) Zero inflation

Given a random vector **n ∈** 𝕀^*N*^ with probability distribution *P* (**n**) and PGF *G*(**z**), consider a new random vector **m ∈** 𝕀^*N*^ with the probability distribution defined as

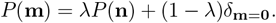

Here *δ*_**m**=**0**_ is the Dirac delta function, which equals 1 if **m** is the all-zero vector **0** = [0, *· · ·*, 0]^⊤^, and 0 otherwise. The PGF of **m** is then given by

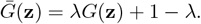

This model is commonly used to describe technical errors encountered during the analysis of single-cell RNA sequencing datasets.

Additionally, the linear mapping approximation (LMA) [25] and its variant [43] allow us to derive PGF solutions for more complex reaction systems using the PGF solutions listed in Table I.

## III. PGF-Based Inference Method for Steady-State Count Data

Building on the PGF solutions of various kinetic models, we first introduce the PGF-based inference method for the steady-state distribution.

Consider a population of *n*_*c*_ cells where the count of the *j*-th species in the *i*-th cell is *n*_*ij*_ for *i* = 1, *· · ·, n*_*c*_ and *j* = 1, *· · ·, N*. Following Eq. (4), the joint empirical PGF (EPGF) for this count data is given by

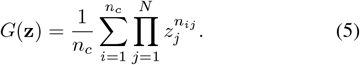

Moreover, from the kinetic model of interest we can derive a PGF, denoted by 𝒢_*θ*_(**z**), where *θ* denotes the kinetic parameters. The inference task is then to estimate *θ* by minimizing the discrepancy between *G*(**z**) and 𝒢_*θ*_(**z**) under a chosen metric. Here, we adopt the mean squared error, defined as

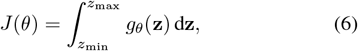

where

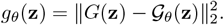

It is worth noting that the mean squared error formulation of *g*_*θ*_(**z**) is a special case of the density power divergence with hyperparameter *α* = 1 (see Eq. (2.1) in Ref. [30]), and that the density power divergence approaches the Kullback– Leibler divergence as *α →* 0 [31]. Here *z*_min_, *z*_max_ ∈ [0, 1] are hyperparameters that define the integration bounds for each component of the PGF vector **z**. The kinetic parameters are estimated by solving the optimization problem

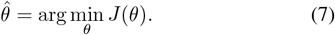

To reduce computational effort, we apply the Gauss quadrature method to approximate the integral Eq. (6) as follows

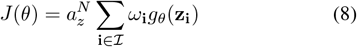

### Algorithm 1

PGF-based inference method for steady-state count data

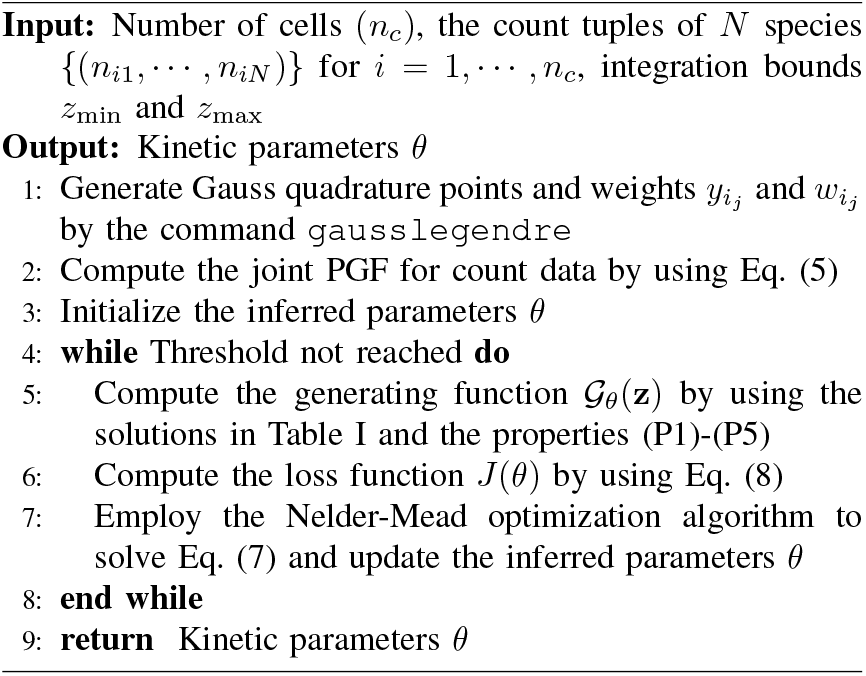

where 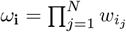,and

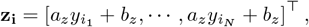

with

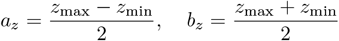

Here 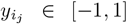 for *j* = 1, *· · ·, N* is the *i*_*j*_-th integration point of the Gauss quadrature of order *N*_*y*_, and 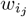 is the corresponding integral weight obtained using the `gausslegendre` function in Julia. The vector **i** = [*i*_1_,, *i*_*N*_]^⊤^ is a sequence of the indices with each component *i*_*j*_ = 1, *· · ·, N*_*y*_ for all *j*, and the set ℐ contains all such index vectors **i**.

The optimization problem in Eq. (7) is solved using the Nelder–Mead algorithm, implemented through the `Optim.jl` package in Julia. Since all kinetic parameters are positive, we play the trick – optimizing their logarithmic transformations and subsequently exponentiating the results to obtain the inferred values. The PGF-based inference procedure is summarized in Algorithm 1.

In Fig. 1a, we illustrate the PGF-based inference method using the telegraph model (inset, Fig. 1a) [44] and its application to single-cell RNA sequencing (scRNA-seq) data. The scRNA-seq data are typically represented as a gene-by-cell count matrix. For a selected gene, we compute the histogram of its transcript counts and, using Eq. (5), convert this histogram into the EPGF. In the telegraph model, a gene switches between active and inactive states with rates *σ*_on_ and *σ*_off_, respectively; transcription occurs only in the active state at rate *ρ*, and mRNA degrades at rate *d*. The corresponding PGF solution 𝒢_*θ*_(*z*) is provided in Table I. The kinetic parameters are *θ* = [*ρ, σ*_on_, *σ*_off_]^⊤^. Under steady-state conditions, the four kinetic parameters cannot be inferred simultaneously; hence, without loss of generality, *d* is set to 1, which is equivalent to normalizing the remaining three parameters by *d*. These parameters are estimated by optimizing the cost function *J*(*θ*) in Eq. (6), where the integral is efficiently evaluated using the Gauss quadrature method (Eq. (8)).

**Fig. 1.**
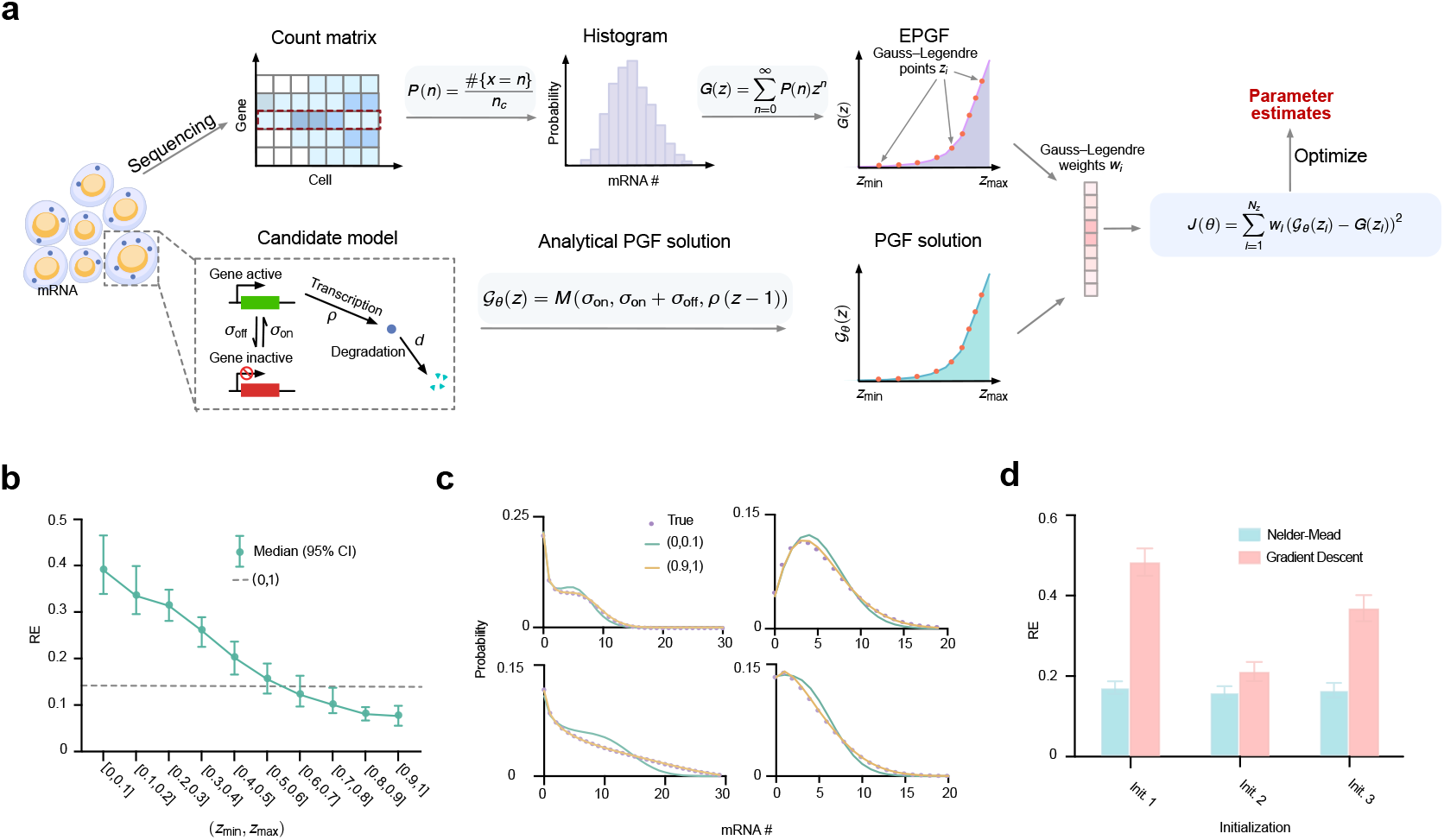
(a) Schematic of the PGF-based inference method using scRNA-seq data and the telegraph model. (b) Inference accuracy over 200 count distributions generated from randomly sampled kinetic parameters increases as the integration range approaches 1. The best accuracy is achieved at [0.9, 1], slightly better than the natural choice [0, 1] (dashed line). Bars indicate the 95% confidence interval of relative errors averaged across all three telegraph model parameters. (c) Reconstructed distributions from four inferred parameter sets using [0.9, 1] (yellow) align more closely with the ground truth (purple dots) than those from [0, 0.1] (green). (d) The Nelder–Mead algorithm outperforms gradient descent and shows robustness to different initialization strategies.

Our PGF-based inference method involves two hyperparameters—the integration bounds *z*_min_ and *z*_max_. To assess their impact on inference accuracy, we uniformly sampled 200 sets of kinetic parameters *ρ* ∈ [1, 30], *σ*_on_ ∈ [0.01, 3], and *σ*_off_ ∈ [0.01, 10]. For each set, we generated steady-state count distributions for 1000 cells using the SSA implemented in `DelaySSAToolkit.jl` [45]. We then performed PGF-based inference with integration ranges [*z*_min_, *z*_max_] varying from [0, 0.1] to [0.9, 1], along with the natural choice [0, 1]. All log-transformed parameters were initialized at 1. As shown in Fig. 1b, the inference accuracy, measured by the relative error averaged over all inferred parameters,

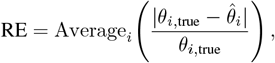

decreases steadily as the integration range approaches 1, reaching its minimum at [0.9, 1], which is slightly smaller than that of the natural choice [0, 1]. To further examine this behavior, we reconstructed count distributions from four different sets of inferred parameters using the integration ranges [0.9, 1] and [0, 0.1]. As shown in Fig. 1c, the reconstructed distributions with [0.9, 1] closely match the ground truth, whereas those with [0, 0.1] poorly capture the distribution tails. This agrees with intuition: according to the definition of the PGF (Eq. (4)), the contribution of each term *P* (*n*)*z*^*n*^ decays rapidly with increasing *n* when *z* is small. Taken together, these results suggest that the interval [0.9, 1] is a practically effective choice for PGF-based inference.

As our PGF-based inference method remains optimization-centered, we next investigate how the choice of optimization algorithm and initialization strategy influences inference accuracy. We consider two optimization algorithms—the Nelder–Mead method and gradient descent, the latter representing a broad class of gradient-based methods—and three initialization strategies: (i) setting all log-transformed parameter values to 1; (ii) using log-transformed MOM estimates (see Appendix A); and (iii) perturbing the log-transformed MOM estimates by adding random values sampled from 𝒩 (1, 1.5). Each algorithm–initialization combination was applied to count distributions generated from 200 sets of kinetic parameters, and the relative error was computed for each case. The results, summarized in Fig. 1c, show that the Nelder–Mead algorithm consistently outperforms gradient descent across all initialization strategies. Moreover, the inference accuracy of Nelder–Mead remains relatively stable across the three strategies, whereas gradient descent exhibits substantial variation, indicating that Nelder–Mead is less sensitive to initialization. We also found that Nelder–Mead requires less computation time than gradient descent, since it is gradient-free and gradient evaluation in our setting involves additional overhead from hypergeometric functions. Taken together, these results suggest that the optimal configuration for the PGF-based inference method is to use the Nelder–Mead algorithm with the simplest initialization strategy—setting all log-transformed parameter values to 1—together with the integration range [0.9, 1].

## IV. Performance evaluation

Given the optimal configuration, we next compare the PGF-based inference method with representatative methods from the other three groups of inference methods metioned in Introduction – ABC, MOM (Appendix A) and MLE integrated with FSP (Appendix B) from the perspectives of accuracy, computational cost and robustness against data contamination.

To this end, we generated five sets of kinetic parameters for the telegraph model (Appendix Table II) and used the SSA to simulate 10 batches of count data for each set, with each batch containing 1000 cells. We first compared the PGF-based inference method with ABC, implemented via `ApproxBayes.jl` using Gamma(2,2) priors and the default error tolerance *ϵ* = 0.1. For each parameter set, both methods were applied to all batches, and the median of RE was computed to obtain a robust estimate of inference accuracy while mitigating random sampling effects. The mean and SEM (standard error of the mean) of these medians are shown in Fig. 2a, demonstrating that the PGF-based method is substantially more accurate than ABC. We also assessed computational efficiency. Both methods were run on a MacBook Air (Apple M2 chip, 16 GB memory), and as shown in Fig. 2b, the PGF-based method was over 500 times faster. Due to this large disparity in speed and accuracy, ABC was excluded from further comparisons. Next, we benchmarked PGF-based inference, MOM, and MLE+FSP across a wide range of sample sizes. Using the same five parameter sets and data generation protocol (with varying sample sizes), we generated count data for comparison. For consistency, all methods employed the Nelder–Mead optimizer with hyperparameters `g_tol` = 10^−20^ and `iterations` = 2000. As shown in Fig. 2c, the averaged RE medians were used to quantify inference error, which decreased with increasing sample size for all methods, as expected. The PGF-based inference method consistently achieved the highest accuracy, with comparable with comparable performance from the others only at very large sample sizes (~ 10^4^). Finally, we evaluated computational time and memory usage (Figs. 2d and 2e). MOM was the most efficient, followed by PGF-based inference, while MLE+FSP was 10–100 times more resource-intensive. Considering both accuracy and efficiency, the PGF-based inference method offers the best balance and is the preferred approach.

**TABLE II.** Parameter values used in Fig. 2.

| Set | $\rho$ | $\sigma_{on}$ | $\sigma_{off}$ |
| --- | --- | --- | --- |
| 1 | 9.64 | 2.76 | 1.66 |
| 2 | 8.76 | 0.97 | 0.19 |
| 3 | 6.26 | 2.86 | 1.00 |
| 4 | 7.24 | 2.32 | 0.80 |
| 5 | 8.54 | 2.60 | 2.69 |

**Fig. 2.**
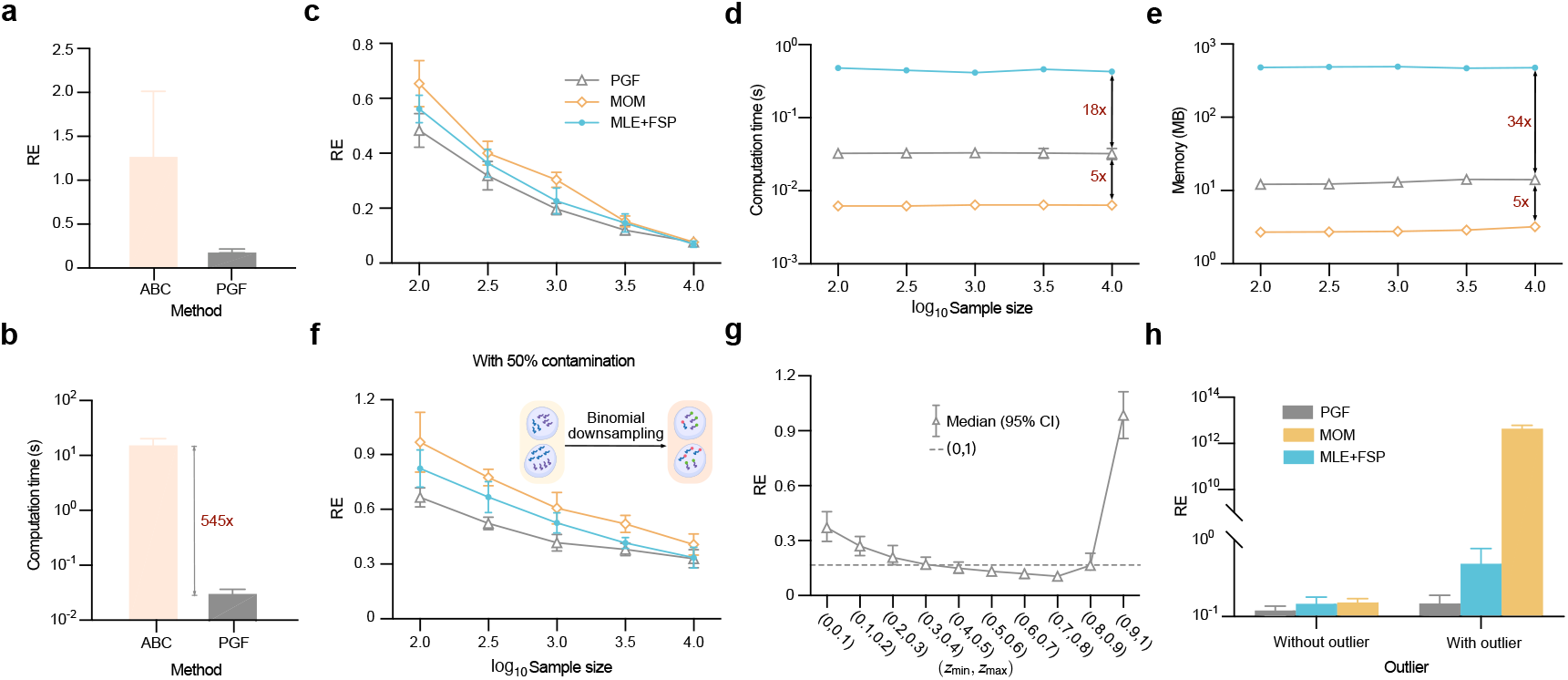
Performance of inference methods in terms of accuracy, efficiency, and robustness. (a) Inference accuracy of PGF-based inference and ABC, evaluated by the mean and SEM (error bars) of median REs across 10 replicate datasets for each of five kinetic parameter sets. (b) Computational time for PGF-based inference and ABC, showing a *>*500-fold speed advantage of the PGF-based method. (c) The mean and SEM (error bars) of median REs as a function of sample size for PGF-based inference, MOM, and MLE+FSP. (d) Runtime and (e) memory usage for the three methods. (f) PGF-based inference remains the most accurate under binomial downsampling (*x*_*i*_ ~Binomial(*n*_*i*_, 0.5)), which mimics sequencing capture inefficiency. (g) Integration range comparison under outlier contamination, showing that [0, 1] achieves the best balance of robustness and accuracy. Error bars indicate the 95% confidence interval of relative errors averaged across all three telegraph model parameters. (h) Inference error under moderate outlier contamination (one count of 30 per batch; sample size = 3,000). PGF-based inference is minimally affected, while MOM shows substantial degradation.

We next evaluated the robustness of the three inference methods by examining how their accuracy degrades under two types of data contamination: binomial downsampling and outliers. The former simulates the sequencing process, where each transcribed mRNA has a probability of being captured and sequenced. This downsampling effect is commonly modeled by a binomial distribution [46]. To assess its impact, we used the same dataset as in Fig. 2c, replacing each count value *n*_*i*_ with a binomial random variable *x*_*i*_ ~ Binomial(*n*_*i*_, 0.5), representing a 50% chance that each transcript is captured. We then applied the same evaluation protocol as in Fig. 2c to compare the three inference methods. As shown in Fig. 2f, although inference accuracy degrades for all methods, the PGF-based inference still outperforms the others, with an even larger performance margin. We also examined robustness to outliers by introducing spurious large values into the data. Specifically, we contaminated the dataset used in Fig. 1b by randomly assigning a count of 100 to one data point per parameter set, simulating extreme measurements. We then followed the same evaluation protocol. As shown in Fig. 2g, under this contamination, the integration range [0.9, 1] is no longer optimal; instead, the natural choice [0, 1] becomes nearly optimal. Taken together with the results in Fig. 1b, these findings indicate that the integration range [0, 1] provides the best balance between accuracy and robustness. Finally, we contaminated the dataset used in Fig. 2c (sample size 3000) by randomly replacing one count per batch with the outlier value 30 and applied the same evaluation protocol. As shown in Fig. 2h, the PGF-based inference method exhibits only a slight increase in inference error, whereas MOM shows a substantial degradation. This confirms that the PGF-based method is the most robust among the three.

In summary, the PGF-based inference method, when combined with the integration range [0, 1], achieves the best overall performance in terms of accuracy, robustness, and computational efficiency (second only to MOM in speed).

## V. Extension to time-resolved count data

Techniques such as single-molecule fluorescent in situ hybridization (smFISH), live-cell imaging, and single-cell EU RNA sequencing (scEU-seq) provide rich time-resolved count data for gene expression dynamics [11], [47], [48], [49]. This motivates an extension of our PGF-based inference method to accommodate time-resolved data. Fortunately, this extension is straightforward to implement. The framework is illustrated in Fig. 3a, using the telegraph model as a representative example. We assume that population-level snapshots of Mrna counts are collected at a set of discrete time points 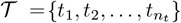. For each time point *t* ∈ 𝒯, we compute the EPGF *G*(*z, t*). In parallel, we evaluate the corresponding analytical PGF solution 𝒢_*θ*_ (*z, t*) from the model at each time point. The discrepancy between the empirical and analytical PGFs is computed analogously to Eq. (8), leading to the following objective function

**Fig. 3.**
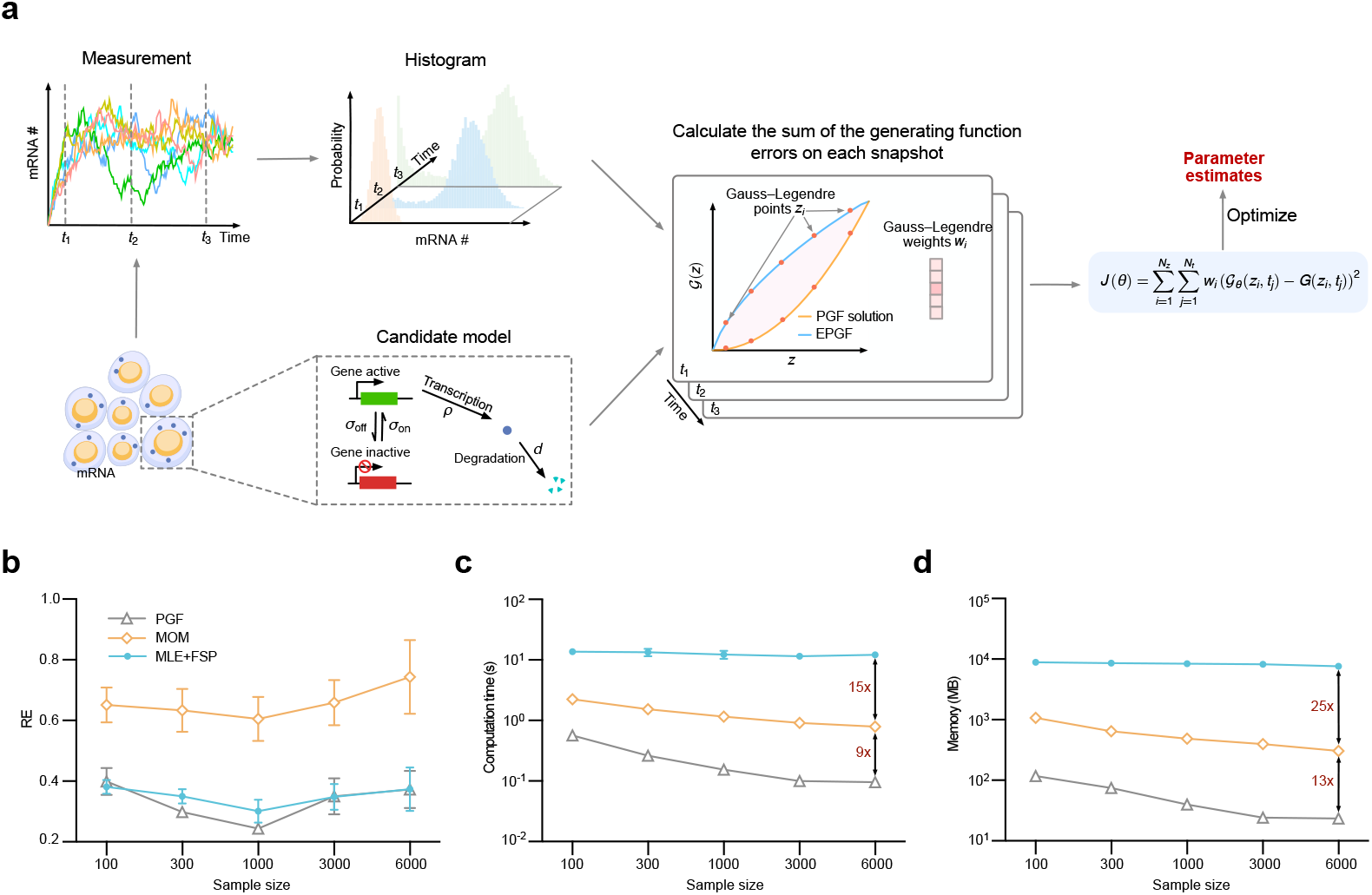
Performance of inference methods on time-resolved count data. (a) Schematic of the PGF-based inference framework applied to time-resolved data. (b) Inference accuracy across varying numbers of cells per snapshot (*n*_*c*_) and time points (*n*_*t*_), with the total number of cells fixed at 12,000. All methods exhibit an optimal trade-off near *n*_*c*_ = 1000 and *n*_*t*_ = 12, with the PGF-based method consistently achieving the highest accuracy. (c) Computational time and (d) memory usage as a function of *n*_*c*_. The PGF-based method is the most efficient, outperforming the other two by one to two orders of magnitude.

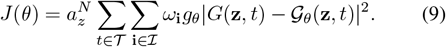

By substituting Eq. (9) for Eq. (8) in Algorithm 1, we obtain a natural extension of the PGF-based inference method for time-resolved count data.

Next, we compared the three inference methods using time-resolved count data. To do so, we reused the kinetic parameters from Fig. 2c and supplemented them with a degradation rate of *d* = 1. Starting from the initial condition where the gene is active and no mRNA is present, we used the SSA to simulate 12,000 cells from time *t* = 0 to *t* = 6. From this simulated dataset, we fixed the total number of sampled cells at 12,000 and varied the number of cells per time snapshot (*n*_*c*_) from 100 to 6,000. The corresponding number of snapshots (*n*_*t*_), evenly spaced over the time interval (0, 6], ranged from 120 to 2. We then followed the same evaluation protocol used in Fig. 2c to compare the three inference methods. Technical details for MOM and MLE+FSP are provided in Appendix A and Appendix B, respectively. To ensure consistency, the optimization hyperparameters were set to `g_tol` = 10^−20^, `f_reltol` = 10^−8^, and `iterations` = 2000. As shown in Fig. 3b, all three methods exhibit an optimal trade-off between temporal resolution (*n*_*t*_) and the number of cells per snapshot (*n*_*c*_), with optimal performance achieved around *n*_*c*_ = 1000 and *n*_*t*_ = 12. Across the entire range of *n*_*c*_, the PGF-based method consistently achieved the highest accuracy. Interestingly, we also quantified the computational time (Fig. 3c) and memory usage (Fig. 3d) for all three methods. In this setting, the PGF-based method emerged as the most computationally efficient—it was an order of magnitude faster than MOM and used only one-tenth of its memory. This improvement arises because, unlike in the steady-state setting where MOM solves only algebraic equations, the time-resolved setting requires MOM to repeatedly solve ODEs for moment trajectories—an overhead that the PGF-based method avoids.

## VI. Model Selection Using PGF-Based Inference for Time-Resolved Count Data

We now describe how to extend the PGF-based inference method for time-resolved count data to address the problem of model selection, with the goal of identifying gene activity dynamics. Since our method does not rely on conventional likelihood functions, classical model selection approaches based on information criteria (e.g., AIC [50], BIC [51]) are not applicable. Instead, we adopt and extend the cross-validation-based strategy proposed in Ref. [32], which was originally developed for steady-state count data.

Assume we collect count data from *n*_*c*_ cells at each time point *t* ∈ 𝒯, where 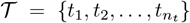. To implement 10-fold cross-validation, we randomly partition the *n*_*c*_ cell-level observations at each time point into 10 equally sized subsamples. For each candidate model, nine subsamples are used to infer the kinetic parameters 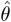, and the remaining subsample serves as validation data, on which the inference accuracy is evaluated via the performance score 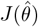 computed from Eq. (9). This process is repeated ten times so that each subsample is used exactly once for validation. The resulting vector of performance scores for each candidate model is denoted by 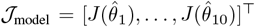. To determine the best-fitting model, we apply the one-standard-error rule [52]. Given a set of competing models {model_1_, …, model_*n*_}, we compute the mean and standard deviation of performance scores for each model

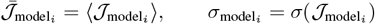

for *i* = 1,, *n*. We identify the model with the lowest mean performance score,

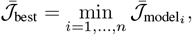

and denote its corresponding standard deviation as *σ*_best_. We then compute the Pearson correlation coefficient between the performance score vector of the best model and that of each candidate model:

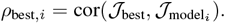

The model-specific performance threshold is defined as

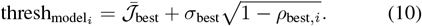

A candidate model is considered competitive if its mean performance score is below this threshold. The full procedure is illustrated in Fig. 4a and detailed in Appendix C.

**Fig. 4.**
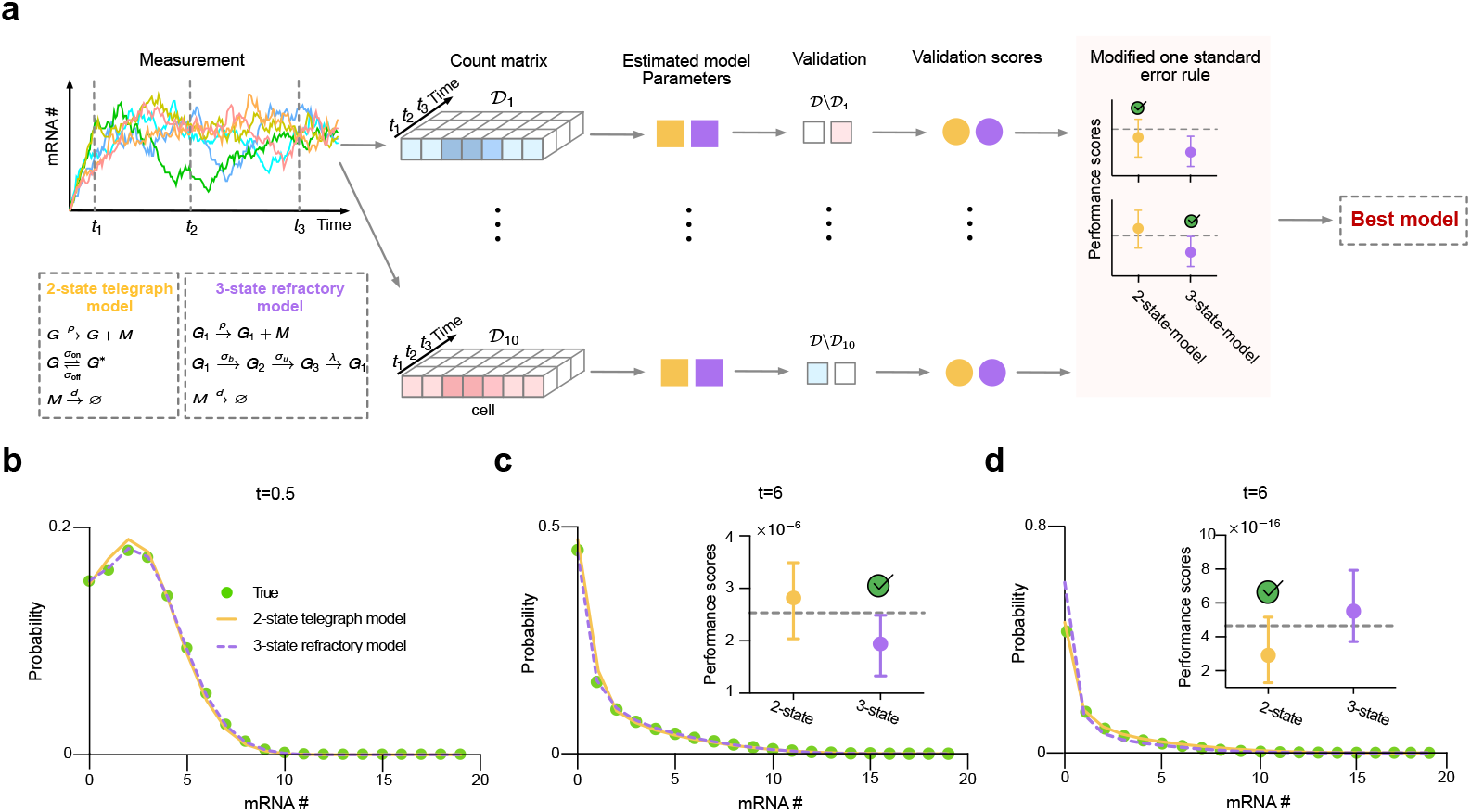
Validation of the PGF-based model selection method using time-resolved count data. (a) Schematic of the PGF-based model selection framework applied to time-resolved data, using the telegraph and refractory models as candidate models (inset). (b)(c) Reconstructed mRNA distributions at *t* = 0.5 and *t* = 6 using inferred parameters from the refractory model (yellow) and the telegraph model (purple) based on one fold of time-resolved data. The refractory model matches the ground truth distribution (green) more closely than the telegraph model. (c, inset) Performance scores across 10 folds identify that the refractory model is correctly selected as the best-fitting model. (d) Using only steady-state data at *t* = 6 results in incorrect selection of the telegraph model (inset). The reconstructed distribution based on the refractory model under this setting fails to capture the ground-truth distribution, particularly at zero-mRNA count.

To validate the proposed model selection method, we considered the refractory model [38], [53], a three-state gene model in which two states are transcriptionally inactive (prohibitive), and the remaining state permits active transcription, as illustrated in the inset of Fig. 4a. Using the kinetic parameters reported in Appendix Table III, we employed the SSA to simulate 1,000 cells from time *t* = 0 to *t* = 6, starting from gene state *G*_1_ and zero mRNA. By *t* = 6, the system reaches steady state. Count data were collected at 0.5 time unit intervals. We evaluated model selection performance by listing the two-state telegraph model as a competing alternative and deriving the time-dependent PGF solution for the refractory model analytically (see Appendix D). Applying the cross-validation–based PGF inference procedure to the time-resolved dataset, the resulting performance scores (Fig. 4c, inset) correctly identified the refractory model as the best-fitting one. This conclusion is further supported by the reconstructed distributions: as shown in Figs. 4b and 4c, the distributions reconstructed from inferred parameters using both the refractory and telegraph models (based on a representative fold; see Appendix Table III) were compared with the ground-truth distribution. The refractory model yields an more accurate match.

**TABLE III.**
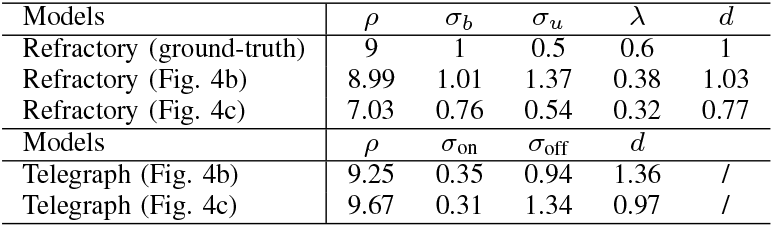
Ground-truth kinetic parameters and parameter estimates for Figs. 4b and 4c.

| Models | $\rho$ | $\sigma_b$ | $\sigma_u$ | $\lambda$ | $d$ |
| --- | --- | --- | --- | --- | --- |
| Refractory (ground-truth) | 9 | 1 | 0.5 | 0.6 | 1 |
| Refractory (Fig. 4b) | 8.99 | 1.01 | 1.37 | 0.38 | 1.03 |
| Refractory (Fig. 4c) | 7.03 | 0.76 | 0.54 | 0.32 | 0.77 |
| Models | $\rho$ | $\sigma_{on}$ | $\sigma_{off}$ | $d$ | |
| Telegraph (Fig. 4b) | 9.25 | 0.35 | 0.94 | 1.36 | / |
| Telegraph (Fig. 4c) | 9.67 | 0.31 | 1.34 | 0.97 | / |

For comparison, we also applied the steady-state model selection method from Ref. [32], using only the snapshot at *t* = 6. In this case, the method incorrectly identified the telegraph model as the best-fitting one (Fig. 4d, inset). The reconstructed distribution from the refractory model under this steady-state-only setting poorly captures the ground-truth, particularly at zero-mRNA levels, suggesting possible overfitting in parameter inference across folds. This is also reflected in the inferred parameter values (Appendix Table III), where estimates based on steady-state data are considerably less accurate than those obtained using time-resolved data—a trend also noted in Ref. [32]. Taken together, these results demonstrate the effectiveness of the PGF-based inference framework combined with cross-validation for model selection using time-resolved count data. Moreover, they highlight the necessity of time-resolved measurements for accurately identifying gene regulatory mechanisms.

## VII. Discussion

In this paper, we extended the PGF-based inference method proposed in Ref. [32]—originally developed for steady-state count data—to accommodate time-resolved count data, and further generalized the associated model selection strategy based on cross-validation. Using this extended framework, we demonstrate that time-resolved data enables the reliable identification of complex gene expression models with multiple gene states, a task that cannot be achieved using traditional steady-state count data alone. In addition, we investigated the effect of key hyperparameters on inference accuracy and identified an optimal configuration for practical use. We systematically evaluated the accuracy, computational efficiency, and robustness under two types of data contamination, of representative methods from four major inference frameworks. Our results show that the PGF-based inference method consistently outperforms the others across nearly all experimental settings and evaluation metrics. These findings highlight the PGF-based approach as a highly promising next-generation inference framework for count data, a common data structure arising in stochastic biochemical reaction systems.

PGF-based inference methods have also been studied in Refs. [30], [31], where inference is performed by minimizing the density power divergence, which involves a hyperparameter *α*. In this work, we similarly experimented with several values of *α* (results not shown), and observed that the inference accuracy was comparable to that obtained using the simpler mean squared error metric, as adopted in Ref. [32] and throughout this paper. It is worth noting that Refs. [30], [31] primarily focused on models with simple analytical PGFs, such as the Poisson and negative binomial distributions. In contrast, the PGFs addressed in Ref. [32] and in the present work arise from biochemical kinetic models and are substantially more complex.

Notably, the PGF-based inference method proposed in Ref. [32] and further developed in the present paper is readily extensible to multi-species biochemical reaction systems. Although this study focuses on the telegraph model involving a single mRNA species - so as to isolate and evaluate inference performance without confounding factors - this extensibility is a key feature for developing kinetic models based on the central dogma of molecular biology. This is particularly important in light of recent advances in single-cell sequencing technologies, which allow for simultaneous measurement of multiple molecular species within the same cell. For example, cellular indexing of transcriptomes and epitopes by sequencing (CITE-seq) enables joint quantification of mRNA and surface proteins [54], single-cell assay for transposase-accessible chromatin using sequencing (scATAC-seq) captures chromatin accessibility alongside transcriptomic data [55], multiplexed error-robust fluorescence in situ hybridization (MERFISH) provides spatially resolved nuclear and cytoplasmic RNA counts [56], and Velocyto extracts spliced and unspliced RNA counts [57]. These developments highlight the importance of modeling frameworks that can flexibly incorporate multiple species.

While the present study focuses on the application of the PGF-based inference method to model selection, future work may explore its integration with other downstream tasks, such as clustering and deconvolution, to further leverage the power of PGF in single-cell data analysis.

## Appendix

### A. MOM-based inference method

One competing approach is the MOM-based inference method, which constructs a synthetic likelihood from the moments of the count data. For clarity, we focus here on the procedure for applying the MOM-based method to infer the kinetic parameters of the telegraph model.

Consider a population of *n*_*c*_ cells, where each cell has *n*_*i*_(*t*_*j*_) molecules of species *X* (e.g., mRNA) measured at time *t*_*j*_, for *j* = 1, 2, …, *n*_*t*_. The first three moments computed from the count data are

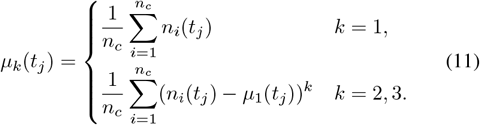

By the law of large numbers, the moment distribution is approximately Gaussian. We use the following likelihood function [10], [12] to infer the kinetic parameters

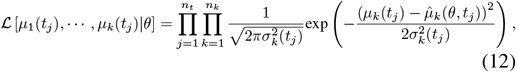

where 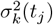 denotes the variance of the *k*-th order empirical moment at time *t*_*j*_, computed from the count data using the following expressions

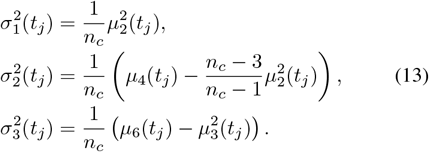

By contrast, the moments 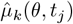 are theoretical moments computed from the underlying kinetic model. For the telegraph model under steady-state conditions, the four kinetic parameters cannot be independently identified; only the ratios of *ρ, σ*_on_, and *σ*_off_ normalized by the degradation rate *d* are identifiable. Therefore, we fix *d* = 1 without loss of generality. In this setting, we set the number of moments *n*_*k*_ = 3 and the number of time points *n*_*t*_ = 1, so that 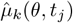 simplifies to 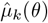. These moments can be directly derived from the steady-state PGF solution provided in Table I, and are given by

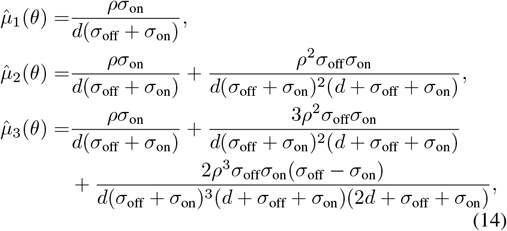

with *d* = 1. Maximizing the likelihood defined in Eq. (12) is equivalent to minimizing its negative logarithmic likelihood, which is given by

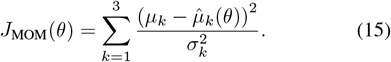

Under steady-state conditions, the numerical procedure for the MOM-based inference method is outlined in Algorithm 2, with optimization details identical to those of PGF-based inference method.

#### Algorithm 2

MOM-based inference method

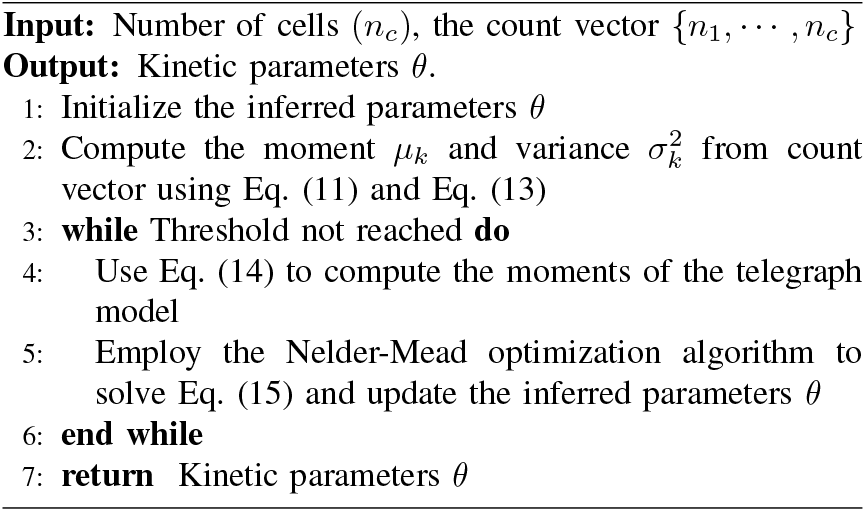

Indeed, Algorithm 2 under the steady state conditions are employed as a parameter initialization strategy in Fig. 1d.

To extend Algorithm 2 to time-resolved count data, we set the number of moments to *n*_*k*_ = 2. In this setting, the reduction in the number of moment measurements is compensated by increased temporal resolution across multiple time points. The number of kinetic parameters to be inferred is four. The theoretical moments 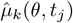 at each time point *t*_*j*_ are computed by solving the system of moment equations

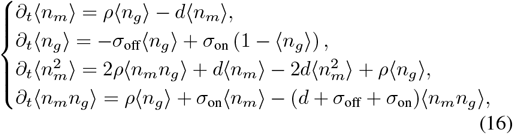

where ⟨·⟩denotes the expected value. Solving this system yields ⟨*n*_*m*_⟩ and 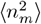 at each time point *t*_*j*_, for *j* = 1, …, *n*_*t*_. These are used to compute the first and second theoretical moments 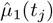 and 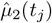. Accordingly, for time-resolved count data, Algorithm 2 is modified as follows: (i) In Step 2, the empirical moments *µ*_*k*_(*t*_*j*_) are computed for *k* = 1, 2 across all time points. (ii) In Step 4, the theoretical moments 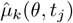 are obtained by numerically solving the moment equations in Eq. (16). (iii) In Step 5, the loss function defined in Eq. (12) becomes

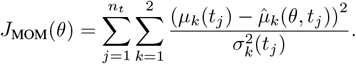

### B. MLE-based inference method

As MLE-based methods are commonly used and serve as natural benchmarks for comparison, we provide the technical details of the MLE-based approach that utilizes the FSP method for likelihood computation.

Given observations from *n*_*c*_ cells measured at time points *t*_*j*_ for *j* = 1, …, *n*_*t*_, the dataset for *N* molecular species is denoted as 𝒟= {(*n*_*i*1_(*t*_*j*_), *· · ·, n*_*iN*_ (*t*_*j*_)) }, where *n*_*ik*_(*t*_*j*_) is the copy number of species *k* in cell *i* at time *t*_*j*_. The total likelihood of observing all data is given by the product over all cells and time points

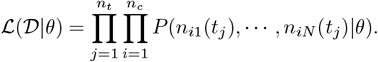

Inference of the kinetic parameters *θ* is then performed by minimizing the negative log-likelihood

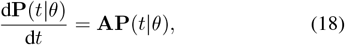

The probability *P* (*n*_*i*1_(*t*_*j*_), *· · ·, n*_*iN*_ (*t*_*j*_) |*θ*) is computed using FSP, which approximates the solution of CMEs by solving a truncated system of ODEs [19]. Specifically, the truncated CME for the telegraph model is given by

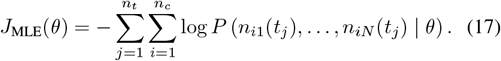

where the probability vector is defined as

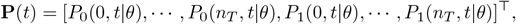

with *P*_*s*_(*n, t* |*θ*) denoting the probability of observing *n* mRNA molecules while the gene is in state *s* ∈ {0, 1} at time *t*, and *n*_*T*_ representing the state space truncation level. The transition rate matrix **A** has the block structure,

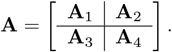

Here the submatrices are given by

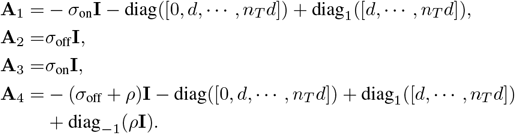

The operator diag_*φ*_(**v**) constructs a diagonal matrix with the elements of the vector **v** placed on the main diagonal when there is no subscript, on the upper off-diagonal when *φ* = 1, and on the lower off-diagonal when *φ* = −1. The identity matrix is denoted as **I**. This system is numerically integrated using standard ODE solvers to evaluate the likelihood required for MLE. Notably, CMEs of any kinetic model can be concisely expressed in the form of Eq. (18) by organizing the probabilities of all possible states into the vector **P**(*t* | *θ*).

The numerical procedure for the MLE-based inference method is outlined in Algorithm 3, with optimization details identical to those of PGF-based inference method.

#### Algorithm 3

MLE-based inference method

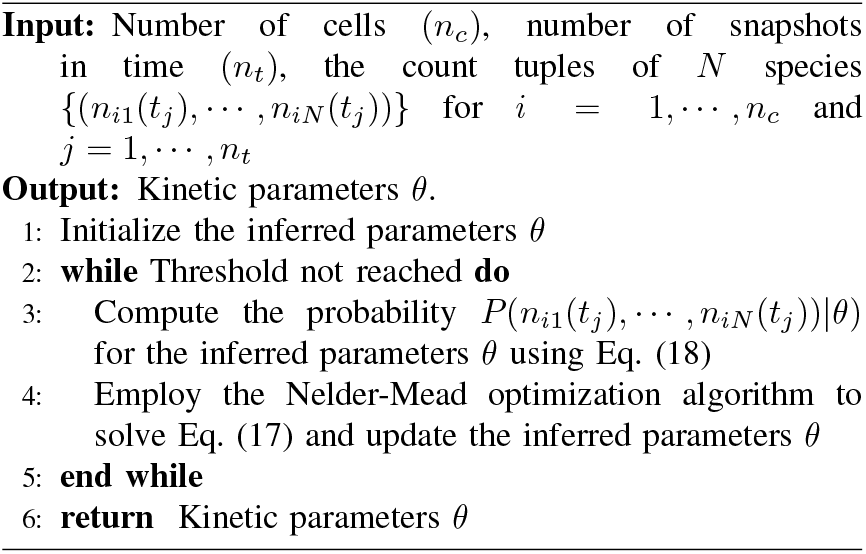

It should be noted that under steady-state conditions (i.e., *n*_*t*_ = 1), there is no need to integrate Eq. (18) over time to obtain the steady-state distribution. Instead, one can directly solve the corresponding stationary system by modifying the equation as follows: replace the first row of the matrix **A** with all ones, and set the left-hand side of Eq. (18) to the vector [1, 0, …, 0]^⊤^. Solving this modified set of algebraic equations yields the steady-state probability *P* (*n*_*i*1_, …, *n*_*iN*_ |*θ*), which is used in Step 3 of Algorithm 3.

### C. Model selection using PGF-based inference method

#### Algorithm 4

Model selection method

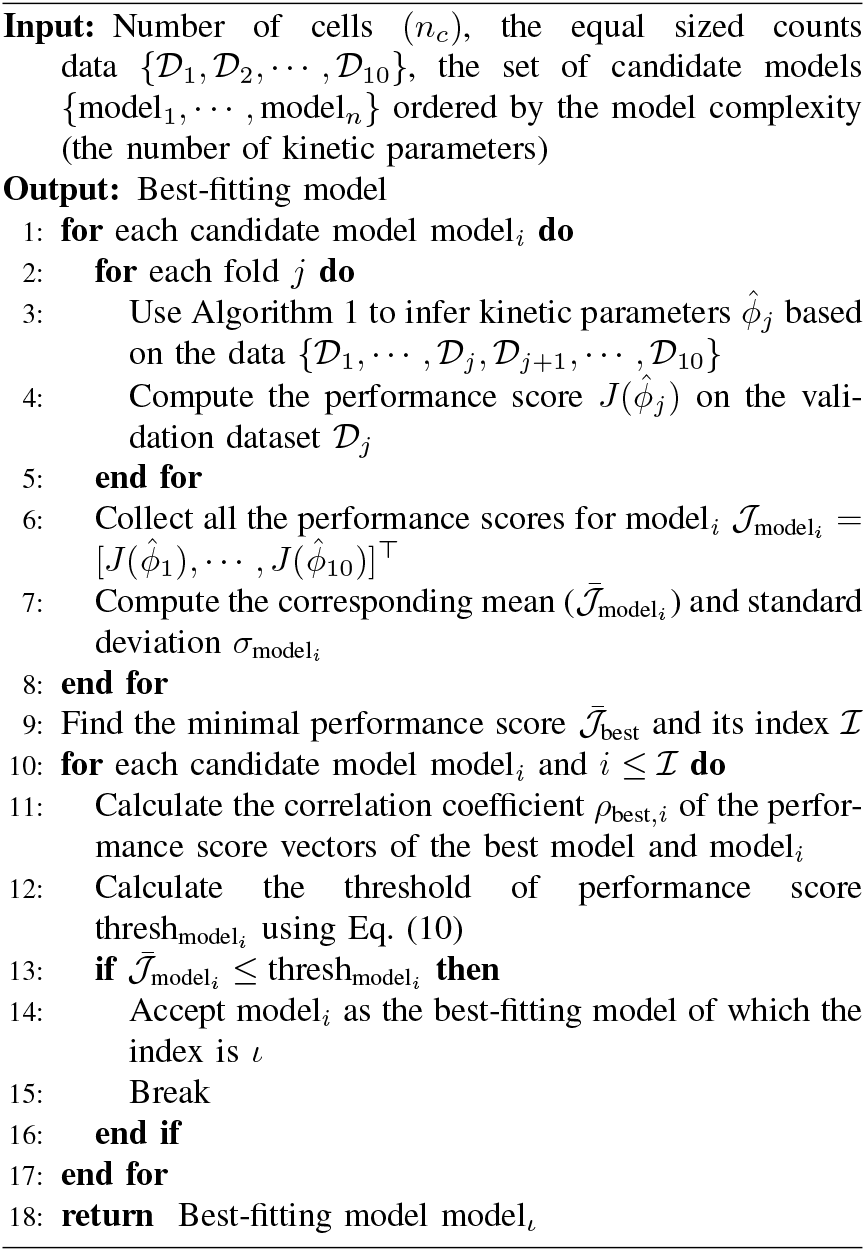

### D. Exact time-dependent solution of the refractory model

For the three-state refractory model illustrated in the inset of Fig. 4a, the corresponding CMEs are given by

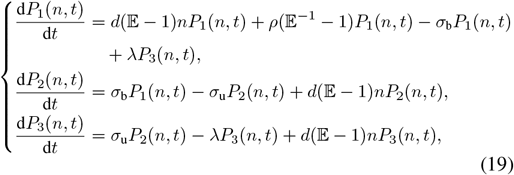

where *P*_*ϕ*_(*n, t*) denotes the probability of observing *n* mRNA molecules when the gene is in state *ϕ* ∈ {1, 2, 3}.

Defining the generating function *G*_*φ*_ = ∑ _*n*_(*u* + 1)^*n*^*P*_*φ*_(*n, t*), Eq. (19) can be recast as the following system of PDEs

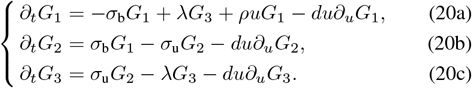

We first use Eq. (20b) to express *G*_2_ in terms of *G*_1_, and subsequently apply Eq. (20c) to represent *G*_3_ in terms of *G*_2_,

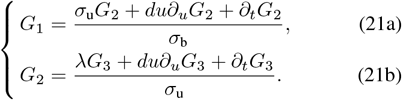

*σ*_u_

Substituting Eq. (21b) into Eq. (22) and simplifying yields an explicit expression for *G*_1_ in terms of *G*_3_,

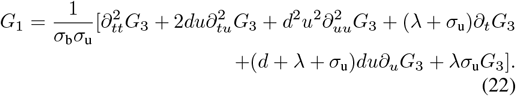

Summing Eqs. (20a)–(20c) and substituting Eq. (22) into the result produces a single parabolic PDE for *G*_3_,

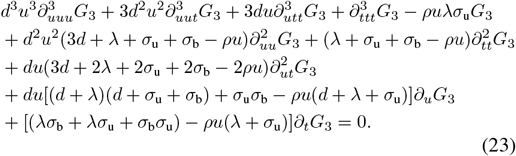

To solve Eq. (23), we introduce the change of variables (*t, u*) ↦ (*w, v*) with *v* = *du* and *w* = ln *du* − *dt*. This transformation can be implemented in Mathematica using the DSolvechangeVariables option. After simplification, the PDE reduces to a third-order ODE,

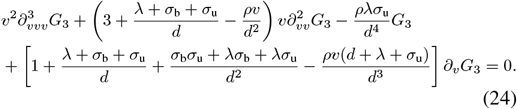

Changing to a description in terms of the variable *x* = *ρv/d*^2^, Eq. (24) further simplifies to

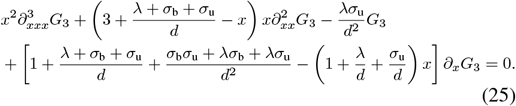

This corresponds to the canonical form of the generalized hypergeometric differential equation (see Eq. (16.8.3) in [58]), from which it follows that the general solution of Eq. (25) is

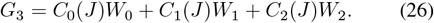

where *J* = exp(*w*). The functions *W*_0_, *W*_1_, and *W*_2_ are defined as follows

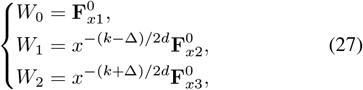

Here *k* = λ + σ_b_ + σ_u_ and 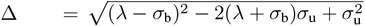. The functions 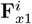 and 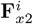 are expressed in terms of generalized hypergeometric functions as

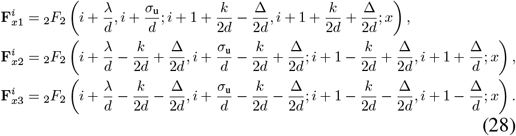

Note that the coefficients *C*_0_, *C*_1_, and *C*_2_ in Eq. (26) are functions of *J* = exp(*w*), and thus depend only on *w*. Their specific forms are determined by the initial conditions.

To this end, we first express *G*_1_ and *G*_2_ in terms of *C*_0_, *C*_1_, *C*_2_, and the basis functions *W*_0_, *W*_1_, and *W*_2_ using Eqs. (22) and (21b). With the substitutions *v* = *du* and *J* = *du* exp(−*dt*), the resulting forms of *G*_1_ and *G*_2_ are

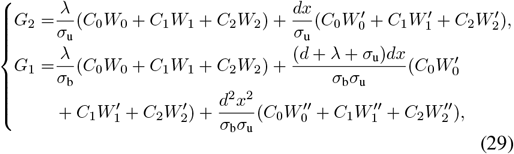

where 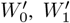, and 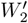 denote the derivatives of *W*_0_, *W*_1_, and *W*_2_ with respect to *x*, given explicitly by

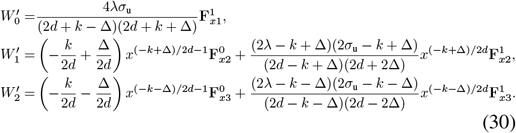

Similarly, 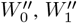, and 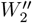 denote the second derivatives of *W*_0_, *W*_1_, and *W*_2_ with respect to *x*, and are given by

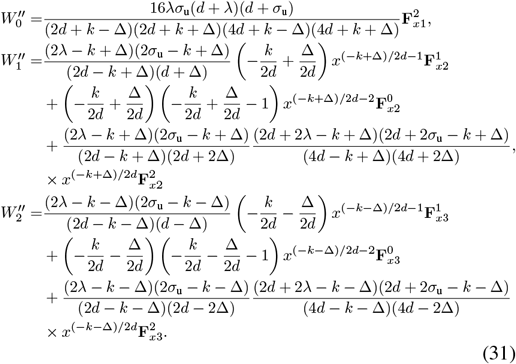

Without loss of generality, we assume that the gene starts in state *G*_1_ with no mRNA transcripts. Equivalently, the initial conditions are *P*_*φ*_(*n*, 0) = 0 for all *φ∈* { 1, 2, 3} and *n ≥* 1 together with *P*_2_(0, 0) = *P*_3_(0, 0) = 0 and *P*_1_(0, 0) = 1. At *t* = 0, we have *J* = *v*. Using the initial conditions together with Eqs. (26) and (29), the coefficients *C*_0_, *C*_1_, and *C*_2_ can be determined from

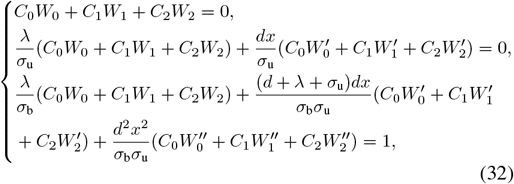

which leads to

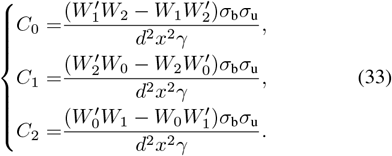

Here

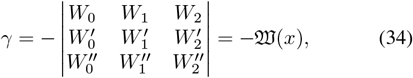

where 𝔚 denotes the Wronskian identity, depending on *W*_0_, *W*_1_, and *W*_2_, and expressed as a function of *x*. According to Abel’s identity [59] and Eq. (25), the following relation can be established for the Wronskian identity,

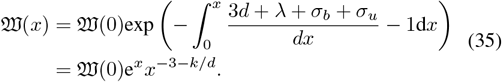

As per the definition of hypergeometric function (see Eq. (16.2.1) in Ref. [58])

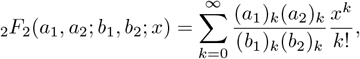

the following limits can be derived

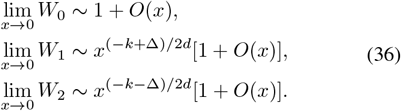

This further leads to

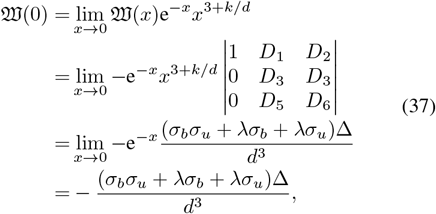

Where

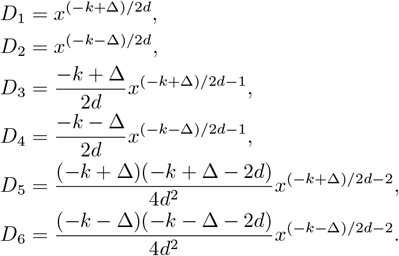

Therefore, it concludes from Eqs. (34), (37) and (35) that

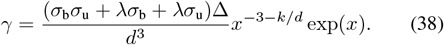

From Eq. (26), the coefficients *C*_0_, *C*_1_, and *C*_2_ in Eq. (33) are functions of the variable *J*, where *J* = *ve*^−*dt*^ reduces to *v* at *t* = 0. Thus, to fully characterize *C*_0_, *C*_1_, and *C*_2_ for arbitrary *t*, we replace *v* with *J*, i.e., *v* ↦ *ve*^−*dt*^ and *x* ↦*xe*^−*dt*^, obtaining

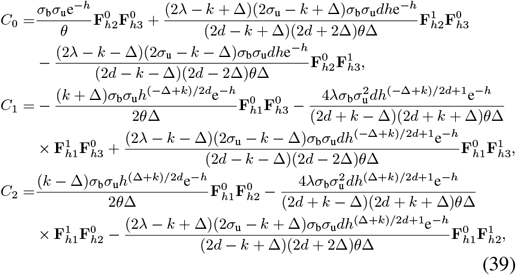

where *h* = *xe*^−*dt*^ and *θ* = *σ*_b_*σ*_u_ + *λσ*_b_ + *λσ*_u_. The functions 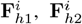, and 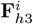 are expressed in terms of generalized hypergeometric functions as

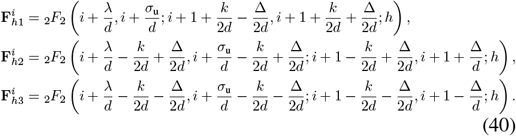

Finally, combining Eqs. (26) and (29), the time-dependent solution of the refractory model, initialized in state *G*_1_ with zero mRNA transcripts, is given by

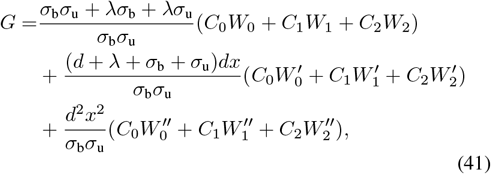

together with Eqs. (27), (30), (31), (39) and (40).

## Notes

This work is supported by NSFC Grants (62573195, 62322309), Shanghai Action Plan for Technological Innovation Grant (23S41900500), and the Natural Science and Engineering Research Council of Canada’s (NSERC’s) Discovery Grant (RGPIN-2024-06015).

### Competing Interest Statement

The authors have declared no competing interest.

